# Fertility intentions and influential factors in Dalian urban city ---A cross-sectional study based on Universal Two-child policy in China

**DOI:** 10.1101/572008

**Authors:** Hongyan Qiu, Qun Zhang, Jin Zhang, Yangjie Ren, Xujuan Zhou, Shuangyue Li, Hong Liang, Jiajia Luo, Qingshan Wang, Liyan Hou

## Abstract

**Background:** In October, 2015, Chinese government announced that one-child policy had finally been replaced by a universal two-child policy. However, the effects of new policy may be far less than expected. So we conducted this research to explore potential influential factors of fertility intention.

**Methods:** This cross-sectional study was conducted and a self-administered questionnaire was designed for collecting socio-demographic information, future fertility intention and influential factors of individual reproductive behavior. The analyses were performed using the SPSS 19.0 statistical software package.

**Results:** A total of 1370 respondents were interviewed. Our research indicated that the mean ideal number of children was only 1.73 and urban respondents’ sex preference was symmetrical preference. 79.1% (884) married people had the first child already, only 7.6% (71) respondents had two children. Among 1370 participants, 30.4% respondents stated that they would have a second child; while 69.6% respondents refused to have two children in future (just wanted only a child). Binary logistic regression analysis (model 1) showed that female, older age, lower education lever, birth place was Dalian, lower family income, the ideal number of children were associated with having 1 child in the future. Model 2 (only respondents with childbearing experience) showed that female, lower family income, couldn’t get additional financial support from parents were more likely refused to have two children; in additional, the ideal number of children and childbearing experience were significantly influences on future fertility intention.

**Conclusion:** Fertility intention and reproductive behavior still below replacement in Dalian city. Our results suggest that several factors (including socioeconomic characteristics, economic factors, desired number of child, childbearing experience) have distinctive effects on fertility intention.

## Introduction

The Chinese government encouraged people to have more children since new China was born in 1949. Due to high fertility and eventual fertility behavior, the population had increased from approximately 6 billion in 1953 to 8.3 billion in 1970[1]. In order to control population growth rapidly, the One Child Family Policy (OCFP) was introduced by central government in China in 1979. In rural areas, OCPC was particularly unpopular because of lacking full pension coverage, sons have traditionally supported parents financially while daughters marring-out of their own families. Due to this reason, from 1984 many provinces allowed couples in rural areas to have a second child if the first had been a girl (Daughter Only Policy). However, urban couples were restricted to a single child. Over the past three decades, the family planning restriction was a “basic national policy” and had curbed the population growth effectively. Since the 1980s, the fertility preference of people throughout China has declined considerably. Total fertility rate dropped from nearly 6 in 1970, through 2.1 in 1991 to 1.22 in 2000[2]. Although there was debate over the true fertility rate for China, data from numerous sources consensus indicated the Chinese fertility has been below replacement level for over 27 years[1, 3].

At least partly as a result of the success of exceptional family planning restrictions in China, population aging is emerging as a major government concern. The percentage of over-60s was 7% of the total population in 2000, extremely rapid reached16.14% in 2015[4]. As the same trends, the percentage of elderly aged 65+ is estimated to increase dramatically from 8.2% in 2010 to 18% in 2030[5]. Rapid population aging will result in a declining labor supply and exacerbating pressure on health systems for the elderly, which would negatively affect development at the macroeconomic level. With a continued ultra-low fertility, the total number of labor force has first declined 3.45 million in 2012 compared to 2011. Now China is facing a new serious demographic crisis that the number of labor force will shrink 8 million per year from 2023[6].

There is no controversy about the neglect effects of current anti-natalist family planning policies as a key factor in preventing China from sustaining the peace of its recent economic growth. So, in November 2013 the Third Plenary Session of the Eleventh Central Committee of the Communist Party of China Changed the anti-natalist family planning policies so that a couple in which one spouse was an only child would be eligible to have a second Child[7]. But by May, 2015, only 1.45 million of 11 million eligible couples applied for permission to have a second child[8]. Several investigations suggested that the effects of new policy may be far less than expected [9–11]. In October, 2015, Chinese government announced that one-child policy had finally been replaced by a universal two-child policy (a couple can have a second child with no restrictions)[12].

China’s population has urbanized rapidly since the “Open Door Policy” in 1978. The one-child rules was strictly enforced for urban couples who in 1980 accounted for about 20% of the population, but 52% in 2012[13]. According to the World Urbanization Prospects data, 70% of China’s population is projected to be urbanized in the 2030[14]. China’s universal two-child policy is highly significant because, for the first time in 36 years, no one in urban city is restricted to having just one child. The fertility intentions and behavior of people who live in cities is likely to create a potentially serious population decline because urbanized rapidly since 1978. People who born between 1980 and 1990 were first generation under the one-child policy have reached the age of childbearing. Additionally, they were first generation to be allowed to have a second child with no restriction.

Although the universal two-child policy has been implemented for three years in China, the birth rate dropped from 12.95‰ in 2016, through 12.43‰ in 2017 to 10.94%o in 2018. Meanwhile the number of newly born babies had decreased from 18.46 million in 2016 to 15.23 million in 2018. Furthermore, the number of newly born babies in 2018 was also lowest since 1961. So we conduct this investigation to examine fertility intentions and future fertility behavior (whether to have two children) of respondents who born between 1980 and 1990 in Dalian city. Additional, we will explore potential influential factors (including socioeconomic status, sex preference, childbearing experiences and external factors such as family support) of fertility preferences using Logistic regression analysis.

## Materials and methods

### Study design and area

We implemented this survey in Dalian, China. Dalian is a large city in northeast China, Which is an important economic center and port city with population approximately 5.9 million. A cross-sectional study was carried out between July and October 2017, using a self-administered questionnaire while investigating people who born between1980 and 1990. To optimize the feasibility of the project, a random sample of women was selected by cluster sampling from 3 of 6 Dalian districts (Zhong shan, Sha he kou and Gan jingzi). The survey areas were randomly selected from the public institutions and companies. A study worksite is defined as a unit within a work unit doing a similar job. Finally, 2 labor-intensive enterprises and 1 technological-intensive factory including 663 subjects, 1 hospital and 10 schools with 707 subjects were recruited in this study.

### Sample size

The sample size was calculated using a formula for cross-sectional studies. The function to estimate the sample size of investigation expressed as following: The sample size was calculated using a formula for cross-sectional studies, n= Z2pq/d2 (α=0.05, d=5%). Our previous survey showed about 30-40% of women refused to have a second child. We assume that the proportion of women refused to have a second child is 40%. In order to estimate the size of sample, we adopt test of two-tailed with p=40%. In addition, 10% of the sample size is increased because of the sub-quality questionnaires; therefore, the total sample size was 1320.

### Data collection

We employed a self-administered questionnaire to obtain data on investigators’ socio-demographic information including birth data, level of education, marital status, and income. In additional, the main contents of the questionnaire include sex preference, childbearing experiences, future fertility intention and influential factors of individual reproductive behavior.

### Statistical analysis

Statistical analyses were performed by SPSS software package version 19.0 (SPSS Inc., Chicago, IL), setting the level of significance at a two-tailed P-value <0.05. Descriptive data are presented as mean with standard error of mean for continuous data and percentage for categorical data.

Categorical variables were performed by Chi-square test. Logistic regression analysis was applied to estimate crude odds ratio (OR) and their 95% confidence intervals (95% CI) between dependent and independent variables. Logistic regression was used to select only those variables which contributed significantly to the variance, with a P-value of > 0.05 as a criterion for removal of a variable from the model.

Family income was classified into 4 categories (<5000, 5000-15000, 10000-15000, and ≥15000); age into 3 (<30, 30-35, and ≥35); birthplace into 2 (Dalian city and other city); and education into 4 (secondary or below, post-secondary degree, college degree, and postgraduate or above); Nature of the worksite into 2 (enterprise, and public institution).

## Results

The total enumerated population was 1370, of which 1070 were women and 300 men. The mean age of these respondents was 32.5±2.95, 25.8% respondents were less than 30years, 48.5% respondents were between 30-35yeas, and 25.7% of respondents were more than 35 years. 63.1% respondents born in Dalian, while 36.9% respondents’ birthplace was other city. 48.4% of respondents were employed by enterprises, 51.6% of respondents were employed by public institutions. Among 1370 participants interviewed, 27.7% were secondary or below, 13.1% were post-secondary degree, 46.1% were college degree, and 13.1% were postgraduate or above. Approximately 80 percent of people had married. A large proportion of respondents had 5000-10000 yuan family income monthly. The detail data were showed in Table 1.

**Table 1.** Demographical characteristics of the participants (n=1370)

| Characteristics | n | % |
| --- | --- | --- |
| Sex |  |  |
| women | 1073 | 78.3 |
| men | 297 | 21.7 |
| Age(years) |  |  |
| $<30$ | 354 | 25.8 |
| 30-35 | 664 | 48.5 |
| $\geq 35$ | 352 | 25.7 |
| Birth place |  |  |
| Dalian | 864 | 63.1 |
| Other city | 506 | 36.9 |
| Education |  |  |
| Secondary or below | 380 | 27.7 |
| Post-secondary degree | 179 | 13.1 |
| College degree | 632 | 46.1 |
| Postgraduate or above | 179 | 13.1 |
| Type of the worksite |  |  |
| enterprise | 663 | 48.4 |
| public institutions | 707 | 51.6 |
| Family income(monthly) |  |  |
| $<5000$ | 266 | 19.4 |
| 5000-10000 | 670 | 48.9 |
| 10000-15000 | 336 | 24.5 |
| ≥15000 | 98 | 7.2 |
| Marital status |  |  |
| unmarried | 260 | 19.0 |
| Married | 1110 | 91.0 |

### The ideal number of children and sex preference

The ideal number of children was collected from the questionnaire and phrased as follow: Without considering other factors, which is the ideal number of children in a family? (Answer: 1, 2, and 3 or more)’. 33.9% respondents stated a preference for only one child, while 59.1% respondents reported the ideal number of children was two children, only 7.0% people reported they want three children. According to our results, the mean ideal number of children was only 1.73. Table 2 shows the ideal number of children desired among respondents according to demographic characteristics. The ideal number of more than one child was more common among those who were born in Dalian (P<0.05). The type of the worksite was another factor that affected a respondent the ideal number of children desired. 67.3% respondents who were employed by public institutions reported they want two children, while only 50.2% respondents who were employed by enterprise stated a preference for two children(P<0.05). Chi-square testes indicated that the educational level and family income were significantly and positively associated with the ideal number of children desired (P<0.05). The results indicated that though women desired on average 2 children than men, but the difference was not significant (p=0.184). There were also no significantly differences in ideal number of children between age, and marital status.

**Table 2.**
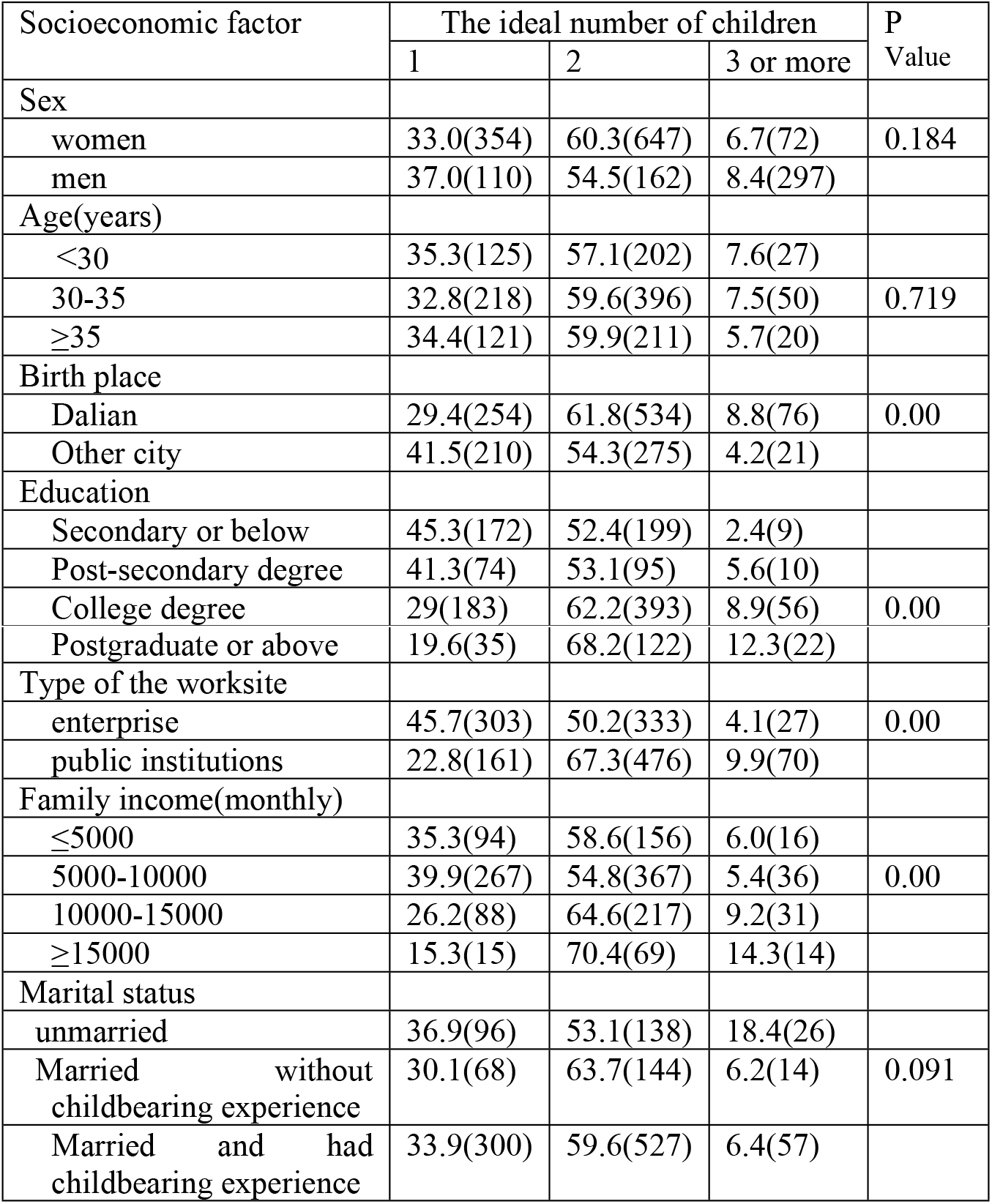
The ideal number of children according to demographic characteristics

The sex preference were collected from 2 questions and phrased as follow:

① The ideal sex composition of offspring (Answer: should have a son, should have a daughter, should have a son and a daughter, don’t care about the sex of child); ② Your attitude about sex composition of your own offspring?(Answer: only boys, only girls, mixed sibset, don’t care about the sex of child).

51.5% respondents stated that ideal sex composition of offspring was a mixed sibset, while 32.9% respondents reported that they didn’t care about the sex of offspring. Interestingly, 11.3% respondents stated a preference for should have a daughter, while only 4.2 respondents reported that ideal sex composition of offspring was should have a boy. Although most respondents thought that ideal sex composition of offspring was a mixed sibset, only 26.2 respondents wanted a mixed sibset; 58% respondents reported that they didn’t care about the sex of own offspring. Similarly to the above results, 11.4% respondents stated just wanted daughters, only 4.4% respondents claimed that they just wanted sons. Our research indicated that urban respondents’ sex preference was symmetrical preference.

### Future fertility prospects (Decision of having one or two child) and influential factors

Approximately 80 percent of people had married. The average age of marriage was 26.7 years, 29.4% people had married before 25 years, and 55.9% had married between 25-30 years, while 14.8% had married later 30years. Among 1370 respondents, 79.1% (884) married people had the first child already, only 7.6% (71) respondents had two children. The women’s mean age at first birth was 28.14±3.40 age. 14.9% people had the first child before 25 years, 57.4% people had first birth between 25-30 years, and 27.5% people had first birth later 30years.

Among 1370 participants, 30.4% respondents stated that they would have a second child; while 69.6% respondents refused to have two children in future (just wanted only a child). Among those who had first child, only 3.2% explicitly stated they would have another child in future, 16.1% respondents might have a second child, 80.7% respondents refused to have two children in future.

Binary logistic regression analysis was applied to estimate the corresponding influential factors of Future fertility (Decision of having two children or just one child). Because after experience the costs of caring for the first child, parents re-assess their fertility intention, so two regression models were generated in order to obtain a better understanding of the determinants of future fertility behavior in Dalian. In effect, the analysis excluded those who never had a child in model 2. Results were adjusted for socioeconomic characteristics, the ideal number of children, and sex preference in model 1, and in model 2, additional adjustment was made for age at first birth, sex of first baby, additional financial support from parents, and childbearing experience.

Model 1 in table 3 shows the corresponding influential factors of Future fertility among 1370 respondents. Compared with men, women were less to have 2 children (P<0.05) (OR=1.55; 95% CI: 1.13-2.14). Age was significantly and negatively associated with fertility intention, respondents who were more than 35 years were more to have only 1 child (P<0.05) (OR=1.87; 95% CI: 1.29-2.73). Education level and family income were significant and negatively associated with having 1 child. Respondents with lower education level or lower family income were more likely to have only 1 child. Decision of having two children or just one child was also significantly influenced by the ideal number of children, respondents’ desired only 1 child were more likely refused to have two children (P<0.05) (OR=7.73; 95% CI: 5.17-11.55). We founded that birth place, type of the worksite and sex preference were no significantly influences on future fertility intention (P>0.05) when the effects of other factors were excluded.

**Table 3.** Adjust Odds ratio with 95% confidence intervals for future fertility prospect and corresponding influential factors

| Characteristics | Model 1<br>OR(95% CI ) | P Value | Model 2<br>OR(95% CI ) | P<br>Value |
| --- | --- | --- | --- | --- |
| Sex |  | 0.007 |  | 0.002 |
| male | 1 |  | 1 |  |
| female | 1.55(1.13-2.14) |  | 2.04(1.31-3.17) |  |
| Age |  | 0.002 |  |  |
| <30 | 1 |  | 1 | 0.143 |
| 30-35 | 1.13(0.83-1.55) |  | 1.00(0.57-1.74) |  |
| ≥35 | 1.87(1.29-2.73) |  | 1.51(0.81-2.83) |  |
| Education |  |  |  |  |
| Postgraduate or above | 1 | 0.005 | 1 | 0.434 |
| Secondary or below | 2.26(1.28-3.98) |  | 1.72(0.72-4.09) |  |
| Post-secondary degree | 1.45(0.67-1.96) |  | 1.21(0.50-2.92) |  |
| College degree | 1.21(0.83-1.77) |  | 0.98(0.54-1.80) |  |
| Birth place |  | 0.079 |  | 0.071 |
| Other city | 1 |  | 1 |  |
| Dalian | 1.55(1.13-2.14) |  | 0.63(0.38-1.04) |  |
| Family income(monthly) |  | 0.000 |  |  |
| ≥15000 | 1 |  | 1 | 0.000 |
| <5000 | 3.30(1.87-5.81) |  | 4.93(2.60-9.36) |  |
| 5000-10000 | 2.55(1.56-4.16) |  | 3.80(1.65-8.70) |  |
| 10000-15000 | 1.67(1.02-2.76) |  | 2.60(1.38-4.88) |  |
| Type of the worksite |  | 0.813 |  |  |
| public institutions | 1 |  | 1 | 0.067 |
| enterprise | 1.05(0.72-1.53) |  | 1.69(0.96-2.95) |  |
| The ideal number of children |  | 0.000 |  | 0.000 |
| 2 children or more | 1 |  | 1 |  |
| 1 child | 7.73(5.17-11.55) |  | 7.60(4.29-13.46) |  |
| Sex preference |  | 0.068 |  | 0.187 |
| No | 1 |  | 1 |  |
| boys | 1.30(0.61-2.78) |  | 0.76(0.25-2.32) |  |
| girls | 0.72(0.37-1.41) |  | 0.50(0.19-1.36) |  |
| mixed | 1.27(0.66-2.45) |  | 0.76(0.28-2.02) |  |
| Age at first childbearing |  |  |  | 0.206 |
| ≥30 |  |  | 1 |  |
| <25 |  |  | 1.25(0.62-2.50) |  |
| 25-30 |  |  | 0.78(0.49-1.23) |  |
| Pressure about home mortgage |  |  |  |  |
| Yes |  |  | 1 | 0.983 |
| No |  |  | 1.00(0.67-1.51) |  |
| Additional financial support from parents |  |  |  | 0.008 |
| Yes |  |  | 1 |  |
| No |  |  | 1.79(1.16-2.76) |  |

Model 2 in table 3 shows the corresponding influential factors of Future fertility among respondents who had childbearing experience (for men whose wife had childbearing experience). According to our results, age, education level, birthplace, type of the worksite, sex preference, age at first childbearing, pressure about home mortagage, parents help to take care of children and sex of first child were no significantly influences on future fertility intention (P>0.05) among respondents who had childbearing experience. Similarly to model 1, women were less to have 2 children compared to men (P<0.05) (OR=2.04; 95% CI: 1.31-3.17); among respondents’ desired only 1 child were more likely refused to have two children (P<0.05) (OR=7.60; 95% CI: 4.29-13.46). Respondents with lower family income were more likely refused to have two children. Furthermore, respondents who couldn’t get additional financial support from parents were more likely to have only 1 child (P<0.05) (OR=1.79; 95% CI: 1.16-2.76). Our results also indicated that childbearing experience was significantly influences on future fertility intention (P<0.05) when the effects of other factors were excluded. Respondents who had painful childbearing experience were more likely to refuse to have two children than those who had happy childbearing experience (OR=7.77; 95% CI: 3.89-15.54).

### Reasons why respondents were unwilling to bear a second child

Due to those respondents who had childbearing experience had lower fertility intention, so we further explored the reasons for not wanting to have two children although Universal Two-child were encouraged by the government (Fig 1). 48.7% indicated that they don’t have enough energy to raise another child, 47% indicated that they couldn’t afford the cost of educating additional children, 27.5% indicated that they couldn’t afford the daily expensive of supporting a second child, 32.3% claimed that one child is enough, 29.8% indicated that poor economic condition couldn’t bring up additional children, 16.9% indicated a fear of older age to have a second child.

**Fig 1.**
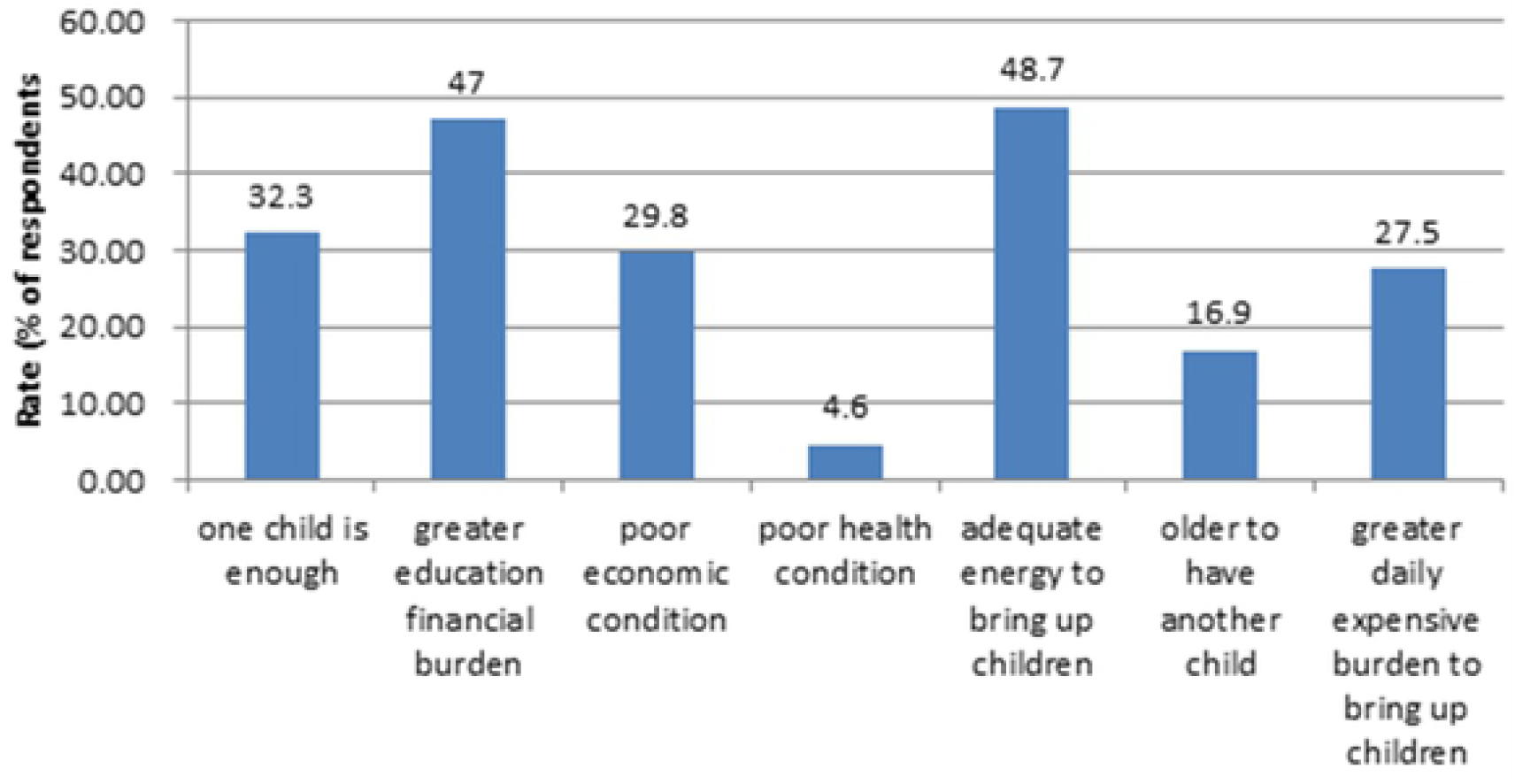
Reasons why repondents were unwilling to bear a second child

## Discussion and Conclusion

Our survey provides results on fertility preferences which support existing evidence of fertility intention still below replacement in Dalian city. Although half of respondents reported that ideal number of children was two children in a family, few respondents wanted more than a total of two children. Approximately 40% of respondents seemed to be an acceptance of the one-child although government announced universal two-child policy.

The mean ideal number of children was only 1.73 among 1370 respondents. Our results were consistent with previous studies in China. For example, Hou reported the mean desired number of children in the period 2000-2010 was 1.5 and 1.82 in urban areas and in rural areas respectively[15]. A survey conducted in Shanghai city indicated that average intended number of children was only 1.1 in 2003[10]. Of note, when above studies were preformed, the ‘one child policy’ was still one of the basic national policies and were strictly enforced among urban areas. Therefore, the ideal number of children was a sensitive word for those respondents and our results might approximate to the actual data.

Most Chinese elderly, especially those in rural areas, lack full pension coverage, so are largely dependent on their children for financial support[16, 17]. Therefore, son preference was prevalent for a long time in rural areas. On the contrary, most urban resident had full pension and medical insurance to support their later life. Our survey indicated that sex preference among Dalian urban residents was balance; most respondents didn’t care about the sex composition of offspring, about a quarter respondents wanted a mixed sibset.

In the last two decades, mean age at first birth was clearly postponed. The fertility rate declines to below replacement level post-1990 in China have been accompanied by increasing ages at first birth [18]. In 2000, the mean age at second birth was 28.5 years[18], however, our results showed that mean age of first birth was 28.14, nearly a third respondents had first birth later 30 years. In China, the average age of marriage and mean age at first childbearing were steadily increasing. Delayed fertility is not an inevitable response to economic and social change. Usually, demands for higher education level, higher income, and own house lead people to delay to have children until they have reached some level of stability. According to a survey conducted in China, decline in mean age at first childbearing would have the opposite effect on the total fertility ratio[18]. Our results also indicated that 16.9% respondents reported a fear of older age to have a second child and nearly half respondents indicated that they don’t have enough energy to raise another child. The postponement of birth has led to low levels of desired and actual fertility in other countries[19, 20]. According to our results, postpone of first marriage and first birth might negatively affect the decision of whether to have a second child in Dalian city.

Modernization and rapid rural-urban migration have been accompanied by reducing fertility intentions in many countries [21, 22]. In contrast to other low-fertility countries, the one-child policy led to a large fall in the total fertility in China. The prime purpose of universal two-child policy was to address the rapid acceleration in population ageing, one of the serious challenges for China in the 21st century. At first, many experts thought that the universal two-child policy is an effective factor that will increase fertility behavior[23]. However, the effects of new policy may be far less than expected. By the end of 2016 (one years later of universal two-child policy), the number of new born babies was only 17.86 million, far lower than expected number of 20-23 million. By the end of 2017 (two years later of universal two-child policy), the number of new born babies was only 17.23 million, even 0.63 million fewer than 2016 in China. The new policy was not resulted in a baby boom as many policy maker expected.

Our results clearly indicated that the role of fertility policy is diminishing fast, and that fertility in China, as elsewhere, is socioeconomically determined. Only 30.4% of respondents would have a second child while most respondents refused to have another child in Dalian. In the Chinese education system, those with high education level earn more money than those with less education. Our results indicated that better-educated respondents and higher family income respondents were both likely to an increased probability of preferring more ideal number of children and having a second child. This is consistent with finding in other researches in China and is explained mainly by socioeconomic and cultural factor. For example, according to a survey conducted in Shanghai city indicated respondents with higher income and education level were more likely to have a second child than those lower income and education level respondents[24]. Another investigation among respondents with one child in urban city also indicated that respondents with higher income were more likely to have additional child than those with lower family income[25]. Economic growth in China was accompanied by inflation and, in urban areas, growing expense of raising children. In additional, high costs of marriage, a raise in housing and increased daily expensive of supporting a family are likely to play an increasingly important role in reducing fertility intentions in China. Our results showed that respondents who couldn’t get additional financial support from parents were more likely to refuse to have a second child. Nearly half respondents indicated that they couldn’t afford the cost of educating additional children, nearly a third indicated that indicated that they couldn’t afford the daily expensive of supporting a second child, 29.8% indicated that poor economic condition couldn’t bring up additional children. Those results all indicated that economic concerns were important factors that reduced fertility intentions in China.

Our investigation indicated that the ideal number of children was important factor that affected a respondent’s decision of having a second child. Compared with respondents with the ideal number of children was more than one child, respondents who stated a preference for only one child were more likely to refuse to have a second child in future. Although continued socioeconomic development and demographic determinants are likely to play an increasingly important role in reducing fertility intentions in China, the other factors of fertility intention are also needed to explore to increase infertility ratio. Our research showed that childbearing experience was major contributing factor of further reproductive plan. Compared with respondents with happy childbearing experience, respondents with painful childbearing experience were more likely to refuse to have two children. After experience of the transition to parenthood parents re-assess decisions about whether to have another child [26]. If having an overall positive experience, or more positive than anticipated, then people should be more likely to have another child. Otherwise, if the transition to parenthood is very difficult or more difficult than expected, then people may refuse to have additional child[27]. Due to strictly one child policy around China, our investigation is the first to assess the relationship between parents’ experience with the first birth and individuals’ further reproductive behavior in China. Both model 1 and model 2 indicated that women were more likely to reject to have two children than men, due to childbirth would further exacerbate gender inequality in China’s workplace, especially for those who take maternity leave. In additional, long conceiving and laboring or complications with cesarean section were challenges for women but not men. May be breast-feeding, depression and sleep deprivation were also important factor for women reject to have additional child. Bianchi and Haas also reported that parenting experiences may be more important for women than men in deciding whether to have another child[28, 29]. Therefore, policy-makers concerned about low fertility should pay much attention to factors that influence the well-being of new parents, and further studies are needed to explore the factors that influence further reproductive behavior of new parents.

A new research reported that low fertility will indeed challenge government programs and very low fertility undermines living standards[30]. 54 governments in 54 countries had enacted policies intended to raise fertility in 2013. In China, more effective policy actions are needed to improve fertility intentions and eventual fertility behavior. The next step, the total removal of the fertility control policy, needs to be considered sooner than later.

## Declarations

### Ethics approval and consent to participate

The study was approved by Ningxia Medical University Ethics Committee. 1370 respondent in the study based on voluntary and provided their informed consent. All respondents are adults. Data were collected by an anonymous, self-administered questionnaire, which was completed by the employees themselves in each study setting. Respondents were advised that no identifying data would be collected, their participation was entirely voluntary and they could withdraw at any time without consequences.

### Availability of data and material

The datasets used and analyzed during current study are available from the corresponding author on reasonable request.

### Funding

This work was supported by National Social Science Foundation of China (16BRK001).

### Authors’ contributions

Hongyan Qiu and Liyan Hou conceived the study and wrote the first draft; Qun Zhang, Jin Zhang and Yangjie Ren contributed data collection; Hong Liang and Jiajia Luo contributed analysis tools; Xujuan Zhou, Huangyue Li and Qingshan Wang revised the manuscript.

## Acknowledgement

We would like to thank Dalian Municipal Center For Disease Control & Prevention for data collection.

## Reference

1. Hesketh T, Zhu WX: The one child family policy: the good, the bad, and the ugly. BMJ 1997, 314(7095): 1685–1687.

2. National Bureau of Statistics of China. Tabulation on the 2000 Population Census of the People’s Republic of China. Beijing: China’s Statistics Press 2002.

3. Zhigang G: Why the fertility-rates in recent years notably ‘pick up’: Evaluation of the 2006 national population and family planning survey. Zhong guo ren kou xue (Chinese Journal of Population Science) 2009, 33:2–15.

4. National Bureau of Statistics of China. Statistical Communique on national economic and social development of People‘s Republic of China in 2012. Beijing: National Bureau of Statistics of China; 2012.

5. Zeng Y WZ: A policy analysis on challenges and opportunities of population/household aging in China. J Popul Aging 2014, 7:255–281.

6. China Statistical Yearbook 2016. Beijing: National Bureau of Statistics of China; 2016.

7. Communique of the Third Plenary Session of the 18th Central Committee of the Communist Party of China. Decision of the Central Committee of the Communist Party of China on Some Major Issues Concerning Comprehensively Deepening the Reform. (2013). 2013.

8. National Health and Family Planning Commission. Press conference of National Health and Family Planning Commission on July 10th, 2015. 2015.

9. Zhihua C HX, Qian C, et al.: Investigation on fertility intention and influencing factors under new fertility policy in Chongqing. Chinese Journal of Family Planning 2014, 22(10):661–665.

10. Rong C BcG: Survey and Research on fertility desire in the past 30 years in Shanghai City. Population and Society 2014, 30(1):49–54.

11. Zhi ke J XtF: Bearing the Second Child Desire of Urban Separate Only Child Couples-Based on the Survey of 558 Youths in 5 Kinds of Industry in Nanjing and Baoding City. Population Journal 2015, 37:5–15.

12. Communique of the fifth Plenary Session of the 18th Central Committee of the Communist Party of China. Beijing 2015.

13. Yongpu Jiang XT: Development of model road of urbanization in China—An Review of recent trends of urbanization. Journal of Guangxi University (Philosophy and Social Science) 2013, 6(35):69–76.

14. Division. UNP: World Urbanization Prospects: The 2014 Revision.; 2014.

15. Hou Jiawei HS, Xin Ziqiang: Transition of Chinese fertility intension:1980-2011. Social Sciences in China 2014, 35(4):78–97.

16. Zhang Y GF: Who will care for the elderly in China?: A review of the problems caused by China's one-child policy and their potential solutions. J Aging Stud 2006, 20:151–164.

17. Weili X zB: Analysis of Influencing Factors of Gender Preference for Endowment in the Countryside. Journal of Northwest A & F University (Social Science Edition) 2015, 2:121–126.

18. Morgan SPZ, G.Hayford, S. R.: China's Below-Replacement Fertility: Recent Trends and Future Prospects. Popul Dev Rev 2009, 35(3):605–629.

19. Myrskylä M, Goldstein, J. R., & Cheng, Y. A.: New cohort fertility forecasts for the developed world:Rise, fall, and reversals. Population and Development Review 2013, 39:31–56.

20. Sobotka T: Is lowest-low fertility in Europe explained by the postponement of childbearing? Population and Development Review 2004, 30:195–220.

21. Assefa N: Fertility is below replacement in Harar Health and Demographic Surveillance System (Harar HDSS), Harar town, Eastern Ethiopia. 2016.

22. Evans L: Italy's fertility rate falls as women reject childbearing. BMJ 1996, 312(7030):530.

23. Yi Zeng TH: The effects of China's universal two-child policy. Lancet 2016, 388(10054):1930–1938.

24. Lin Li YC: Analysis of Influencing Factors about having two children among married people in Shanghai city. Reproduction and Contraceptio 2014, 34(11):914–919.

25. Chen Xh: An investigation on the influencing factors of second child fertility willingness among respondents aged 30-38 in urban city. Yiyao Qianyan 2017, 7(2):304–305.

26. Campbell EK, Campbell PG: Family size and sex preferences and eventual fertility in Botswana. J Biosoc Sci 1997, 29(2):191–204.

27. Margolis R: Parental Well-being Surrounding First Birth as a Determinant of Further Parity Progression. 2015.

28. Haas L: Parental leave and gender equality: Lessons from the European Union. Review of Policy Research 2003, 20:89–114.

29. Bianchi SM, Milkie, M. A., Sayer, L. C., & Robinson, J. P.: Is anyone doing the housework? Trends in the gender division of household labor. Social Forces 2000, 79:191–228.

30. Lee RM, A.: Is low fertility really a problem? Population aging, dependency, and consumption. Science 2014, 346(6206):229–234.

